# Integrative structural modelling of macromolecular complexes using Assembline

**DOI:** 10.1101/2021.04.06.438590

**Authors:** Vasileios Rantos, Kai Karius, Jan Kosinski

## Abstract

Integrative modelling enables structure determination of macromolecular complexes by combining data from multiple experimental sources such as X-ray crystallography, electron microscopy (EM), or crosslinking mass spectrometry (XL-MS). It is particularly useful for complexes not amenable to high-resolution EM—complexes that are flexible, heterogenous, or imaged in cells with cryo-electron tomography. We have recently developed an integrative modelling protocol that allowed us to model multi-megadalton complexes as large as the nuclear pore complex. Here, we describe the Assembline software package, which combines multiple programs and libraries with our own algorithms in a streamlined modelling pipeline. Assembline builds ensembles of models satisfying data from atomic structures or homology models, EM maps and other experimental data, and provides tools for their analysis. Comparing to other methods, Assembline enables efficient sampling of conformational space through a multi-step procedure, provides new modeling restraints, and includes a unique configuration system for setting up the modelling project. Our protocol achieves exhaustive sampling in less than 100 – 1,000 CPU-hours even for complexes in the megadalton range. For larger complexes, resources available in institutional or public computer clusters are needed and sufficient to run the protocol. We also provide step-by-step instructions for preparing the input, running the core modelling steps, and assessing modelling performance at any stage.

## Introduction

Macromolecular complexes are crucial to many biological processes. The function of complexes depends on their three-dimensional (3D) structure–the relative arrangement of the subunits, which assemble to form structural scaffolds, active and ligand binding sites, and regulatory modules^1^. Determining the structure of complexes is therefore key to understand how they assemble and function.

One of the most widely used methods for determining structures of large macromolecular complexes is cryo-electron microscopy (cryo-EM). It involves a 3D reconstruction of a so-called EM density map, which is used to build a structural model. The resolution of cryo-EM maps is sometimes sufficient to build an atomic model using only information of the map. Frequently, however, the high resolution is limited to more rigid regions of a complex, while flexible peripheral domains, that are less well-resolved, remain ambiguous^2, 3^. Moreover, some complexes are difficult to resolve at high resolution due to technical challenges specific to the given sample or sample heterogeneity^4^, e.g. cell extracts containing multiple complexes^5^. Finally, recent work has demonstrated that EM maps of individual complexes can be obtained in their native cellular environment by applying cryo-electron tomography (cryo-ET) to vitrified cells^6–10^, followed by sub-tomogram averaging^11^. In-cell cryo-ET foreshadows a new era of native structural biology, but currently the cryo-ET maps rarely reach resolution beyond 1 nm^8, 12^. Thus, methods for interpreting low-resolution EM maps will be crucial for realizing the potential of in-cell structural biology.

Integrative structural modelling is a method that allows determining structures based on low-resolution EM maps^13^, or even when an EM map cannot be obtained^14–16^. It leverages information from other structural biology techniques such as X-ray crystallography, homology modelling, nuclear magnetic resonance (NMR), small-angle scattering (SAXS), and crosslinking mass spectrometry (XL-MS)^17, 18^. Integrating data from different techniques allows building models at a higher precision than using a single technique alone^8, 19, 20^. Multiple structures of macromolecular complexes have been obtained this way, even under in-cell conditions^7, 8, 21^.

In a typical integrative modelling workflow^19, 22^, structures of individual subunits or domains are first collected using X-ray crystallography, NMR or homology modelling^23^. In some modelling software, the structures are converted to a coarse-grained representation. Second, the experimental data are translated into spatial restraints and used to define a scoring function for subsequent optimization. Third, various optimization methods are applied to find an arrangement of the input structures which minimizes the scoring function. The output is not a single model, but rather an ensemble of models equally satisfying the experimental restraints. Finally, the precision and exhaustiveness of sampling are assessed^20^ to interpret the models and their uncertainty.

We have developed a versatile integrative modelling protocol that can be applied to complexes as large as 50 – 100 MDa nuclear pore complexes (NPCs)^7, 21, 24^, using high- and low-resolution EM maps derived from cryo- and negative stain EM, and cryo-ET maps resolved in cells. The protocol is a computational pipeline integrating our Xlink Analyzer^25^ graphical interface for input preparation, UCSF Chimera software^26^ for EM fitting and analysis of results, and the programming libraries of Integrative Modeling Platform^22, 27^ (IMP) and Python Modeling Interface^28^ (PMI; an interface to IMP by the same authors) for modelling. The protocol implements a multi-step scoring and sampling procedure that enables efficient exploration of conformational space. In addition to the programming interface of IMP and PMI, our protocol offers a straightforward procedure to initially define the target system through the graphical user interface (GUI) of Xlink Analyzer^25^ and text-based configuration files, implements additional restraints, and provides command-line scripts that can be applied to modelling cases without any modification by the user. Here, we present the details of the protocol, installation, and setup instructions for the Assembline software package, which implements the protocol (Supplementary Manual). We also provide step-by-step guidelines for applying it to any protein complex.

## Development of the protocol

We developed the first version of our protocol to build a model of the scaffold of the human nuclear pore complex based on cryo-ET and XL-MS data^24^. Subsequently, we used it to model a yeast Elongator complex based on negative-stain EM maps and XL-MS data^29^, which was later validated by a high-resolution cryo-EM structure^30^. Recently, we have used an updated version of the protocol to build models of yeast NPCs^7, 21^, and to model peripheral subunits of the mycobacterial Type VII secretion system^31^. In this work we are describing the most recent version of the protocol as used for the *Saccharomyces cerevisiae* nuclear pore complex^7^ (ScNPC), now organized in a software package called Assembline.

### Availability

The Assembline software is freely available as an open-source Python package (website: https://www.embl-hamburg.de/Assembline/, git repository with code: https://assembline.readthedocs.io/en/latest/<u>#</u>), which can be installed from source code or from the Anaconda repository (https://anaconda.org/kosinskilab/assembline). Documentation is provided in the Supplementary Manual and can also be found online (https://assembline.readthedocs.io/en/latest/), and in the form of the step-by-step tutorials for modelling the yeast nuclear pore complex (Supplementary Tutorial 1, online version: https://scnpc-tutorial.readthedocs.io/en/latest/) and Elongator complex (Supplementary Tutorial 2, online version: https://elongator-tutorial.readthedocs.io/en/latest/). All the data sets needed for the step-by-step tutorials (i.e., for yeast NPC and Elongator complex modelling) are provided in https://git.embl.de/rantos/scnpc_tutorial.git (for the yeast NPC) and https://git.embl.de/kosinski/elongator_tutorial.git (for the Elongator complex).

### Applications

Assembline can model protein complexes based on EM, XL-MS and protein-protein interaction data, e.g., affinity pull-down experiments indicating protein or domains interactions. Assembline supports other experimental data types through restraints available in a programming interface from IMP and PMI. When only EM data is available, Assembline can be used as a fitting program for fitting multiple structures simultaneously. Although in our published applications we have primarily used EM data, Assembline can be also applied to cases where EM data is not available, for example to build approximate topological models based on XL-MS data only. Complexes from simple dimers to assemblies of multiple subunits and complex symmetries are also amenable. For highly symmetric complexes, Assembline can model complexes with hundreds of subunits, as demonstrated by our applications to NPCs or bacterial secretion systems.

Exemplary results from our previous applications are described in the **Steps 1-6,** installation, set up of virtual environment for modelling and project directory architecture: ∼2 h per modelling project, assuming the input datasets are already prepared.

Steps 7-10, calculation of fit libraries: ∼1 h - 5 h, depending on number of input PDB structures to be fitted, number of EM maps to be used for fitting and allocated computational resources. The calculation of fit libraries for ScNPC takes roughly 3-5 h per input structure and for Elongator complex takes around 1-3 h per input structure.

Steps 11-31, modelling with global optimization and analysis pipeline: ∼1 h to days, depending on modelling parameters, system configuration settings, size of the complex, and allocated computational resources. A single global optimization run (using a single CPU core) for ScNPC takes around 1 h (approximately 20,000 runs to be performed) and for Elongator complex - 6 min (approximately 1,000 runs to be performed).

Steps 32-34, modelling with the recombination step and analysis pipeline: ∼1 h to days, depending on modelling parameters, system configuration settings, size of the complex, the number of selected models to be used for recombination, and allocated computational resources. A single recombination run (using a single CPU core) for Elongator complex takes around 4 min (approximately 1,000 runs to be performed).

Steps 35-40, modelling with refinement and analysis pipeline: ∼3 h to days, depending on modelling parameters, system configuration settings, size of the complex, the number of selected models to be used for refinement, and allocated computational resources. A single refinement run (using a single CPU core) for ScNPC takes around 3-4 h (approximately 20,000 runs to be performed) and for Elongator complex takes around 20 min (approximately 1,000 runs to be performed).

Anticipated results section.

## Overview of the procedure

In this section, we describe the algorithmic details of Assembline (Fig 1). The exact step-by-step procedure is presented in the **Procedure** section.

**Figure 1.**
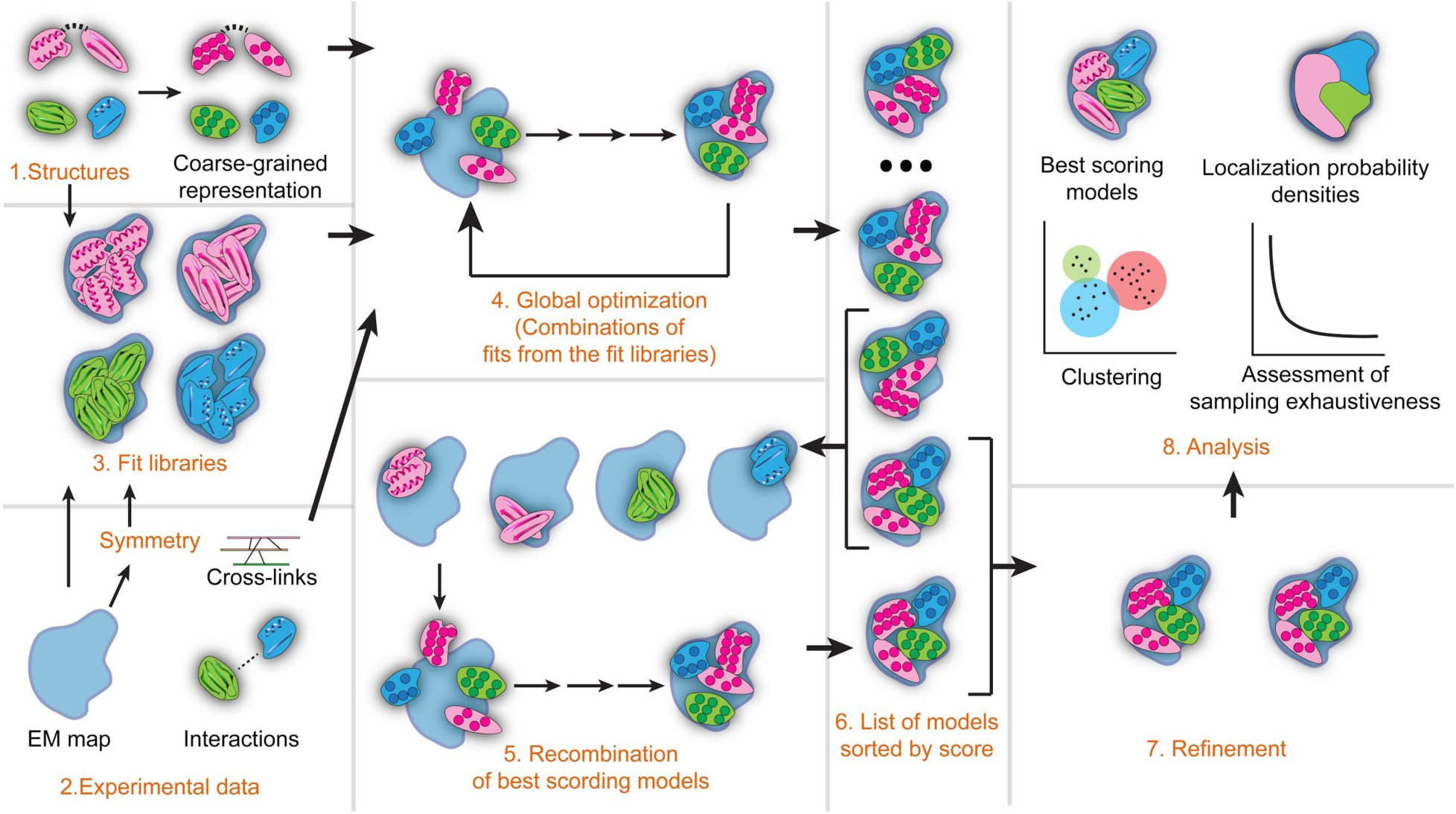
Assembline workflow. Available structures and experimental data are specified as input through configuration files. Optionally, input atomic structures are converted to coarse-grained beads (steps 1 and 2). If EM data is used, libraries of fits for each structure can be calculated (step 3). The fits of all structures are sampled simultaneously by combining fits from the libraries and scored based on input restraints (global optimization, step 4). The best scoring models from the global optimization can be additionally recombined, generating potentially better scoring models (step 5). The best scoring models (step 6) are selected for local rigid-body or flexible refinement (step 7). During analysis, sampling convergence and exhaustiveness are assessed and the output integrative models are analyzed with respect to restraint satisfaction (step 8).

The core algorithmic feature of our protocol is a unique multi-step sampling procedure that efficiently explores structural configurations when an EM map is available. Modelling protein complexes based on EM data, particularly at low resolution, is computationally demanding due to the need of sampling a large conformational space (locations and orientations of multiple subunits within the map) and costly calculations of the cross-correlation between the EM map and the modelled structure. Assembline overcomes both challenges through a global optimization step that first calculates ensembles of fits of individual subunit and domain structures (called “fit libraries”) and then generates good scoring combinations of those fits. Since the scores of the fits have been calculated *a priori*, there is no need to re-calculate the cross-correlation during the optimization, drastically speeding up calculations and enabling efficient sampling of the conformational space. Notably, as the original EM scores were calculated using the original atomic structures, the EM scores are derived from the atomic representation even if proteins are coarse-grained for the actual optimization. The models from the global optimization can be then used as input in the next steps, to either enrich conformational sampling through a recombination between top scoring models and/or refine the complexes using a conventional sampling, in which rigid bodies are moved in a continuous space through random rotations and translations and cross-correlation is calculated “on-the-fly”.

### Representation of proteins for modelling

During modelling, input protein structures can be represented in either atomic or, to accelerate computation, coarse-grained representation as beads at a desired “resolution” (e.g., one bead represents ten amino acid residues). As a trade-off between computing efficiency and accuracy, multiple coarse-grained representations can be used in parallel according to the precision of restraints. For example, a ten-residue per bead representation can be used for the costly excluded volume restraint (steric clash score), while the C_α_ representation can be used for XL-MS data-based distance restraints. The multi-scale representation is implemented using the PMI library^28^.

Although structures are represented as rigid bodies, conformational flexibility can be implemented in two ways. First, flexible linkers can be added explicitly as single-residue beads connected by distance restraints. Second, the input structures can be divided into smaller rigid bodies and restrained using an elastic network to prevent distortions of the input structure.

Additionally, homo-oligomeric complexes and cyclic symmetry are fully supported. Assembline automatically creates the copies of the input structures and imposes symmetry based on user-provided symmetry definitions, and resolves an ambiguity arising when restraints are applied to multiple copies of the same subunit. For example, for crosslink or interaction restraints, it is sufficient that only one of the copies satisfies the restraint.

### Scoring function

The scoring function for modelling is a linear combination of restraint scores as implemented in IMP^27^. Several custom restraints have been implemented along with restraints directly sourced from IMP. Custom restraints in Assembline include: EM restraints based on the *p*-values of fits in the fit libraries (see **Calculation of fit libraries**), symmetry restraints, excluded EM densities restraints—which can be used to penalize penetration of certain EM densities e.g., a segmented membrane density, EM density proximity restraints (e.g., to favor interactions of membrane-binding proteins with a membrane), crosslinking restraints including a log-harmonic crosslink restraint^14^, elastic network restraints for preservation of protein complex interfaces, and binary inter-protein or inter-domain interactions. Standard restraints sourced from IMP include connectivity distance between neighboring domains in sequence, a steric clash score, standard EM fit restraint based on cross-correlation, and EM envelope penetration restraints.

### Calculation of fit libraries

The first step of our protocol is the generation of fit libraries, which are ensembles of non-redundant rigid body fits of the input atomic structures within the provided EM maps (before coarse-graining). The fit library provides alternative positions during the subsequent global optimization step. Typically, hundreds to thousands of fits are generated per structure. If the initial model of the complex has already been constructed by other means, or no EM map is used as a restraint, this step can be omitted and the model can be optimized immediately by the refinement step.

Fit libraries are generated using the FitMap tool from UCSF Chimera^26^ in “global search” mode. A command line interface is provided to fit multiple structures into multiple maps (for example different versions of the EM map or maps of different states of the protein complex). The resulting fits are then clustered using the built-in clustering feature of UCSF Chimera FitMap, wherein representative models from each cluster constitute the final ensemble of non-redundant fits. The user can modify several parameters including the number of random starting positions for the global fitting, clustering threshold settings, resolution of simulated densities and three different cross-correlation score types provided by UCSF Chimera (see the documentation of UCSF Chimera for details https://www.cgl.ucsf.edu/chimera/current/docs/UsersGuide/midas/fitmap.html).

Finally, *p*-values are calculated from the cross-correlation scores of alternative fits as published before^7, 21, 24, 29, 31, 32^. Briefly, the cross-correlation scores are first transformed to z-scores (Fisher’s z-transform^33^) and centered, from which two-sided *p*-values are computed using standard deviation derived from an empirical null distribution (derived from all obtained unique fits and fitted using fdrtool R-package^34^). All *p*-values are then corrected for multiple testing using Benjamini–Hochberg procedure^35^. The corrected *p*-values are used as restraints during the global optimization step. The option of reconstructing the best-scoring fitted models at atomic representation for visual inspection is also available. In addition, the user can use *p*-values to identify unambiguous fits that can be kept fixed in their positions in the subsequent modelling steps to limit the conformational search space.

### Global optimization

The objective of this step is to generate models of the entire complex based on the fit libraries via simultaneous sampling of alternative fits using Monte Carlo simulated annealing optimization^36^ (Figure 1 step 4). The optimization algorithm randomly draws upon the pre-calculated fits from the libraries to generate candidate combinations, scores the combinations according to their fit in the EM map and satisfaction of other restraints, and iteratively repeats the random fit selection and scoring to find better scoring solutions. The EM fit restraint is calculated based on *p*-values pre-calculated during the previous stage, and as such is derived from the atomic representation. Usually, this step consists of hundreds to thousands of independent runs (i.e., optimization trajectories restarted from random initial orientation), each run includes thousands of Monte Carlo sampling steps, depending on the size of the system. The scoring function, representation of structures, and optimization algorithms are implemented using functionalities from the underlying IMP and PMI programming libraries.

This step outputs an ensemble of alternative integrative models, each scored based on violation of the restraints comprising the scoring function. At this point, the user can assess the sampling performance and select final models (see **Analysis** section) and/or continue to the next integrative modelling stages.

### Recombination

The optional recombination stage (Figure 1 step 5) allows enriching the sampling of good-scoring models. This is particularly useful for systems with many rigid bodies or thousands of alternative fits in the fit libraries. In this step, the global optimization protocol is run again, but this time using only the pre-calculated alternative fits of the subunits that led to top models in the global optimization run. The pre-calculated fits that led to those models are retrieved from the original libraries to create smaller fit libraries, and used as input to the optimization algorithm identical to the global optimization. As the fits are coming from good scoring models, this leads to preferential sampling of the conformational space in best-scoring areas and often yields models with scores significantly better than in the first global optimization stage.

### Refinement

Refinement (Figure 1 step 7) optimizes integrative models from the global optimization (including the recombination) step using the underlying IMP and PMI programming libraries. The main difference between the global optimization and refinement is that in the global optimization the EM fit restraint is pre-calculated from *p*-values of fits in the fit libraries, while in the refinement the raw cross-correlation coefficient is used and calculated “on the fly”. This difference is based on the fact that the component (i.e., rigid body) positions in the refinement stage are not drawn from the ensemble of pre-calculated fits but translated and rotated in small increments.

In practice, an ensemble or single best scoring model(s) can be used as input. The refinement calculations can be computationally expensive. Therefore, it is recommended for the user to provide input structures that are already approximately fitted to the EM map (e.g., output structures from the global optimization). Refinement can also be used independently of any prior integrative modelling runs, e.g., to optimize models obtained using other modelling software.

The refinement, as in the case of global optimization, starts with representing the defined system at desired coarse-grained resolutions and proceeds with stochastic sampling of alternative rigid body conformations by Monte Carlo simulated annealing. The coordinates of flexible beads are optimized using the conjugate gradient algorithm. All output models are sorted by total score and, at this point, the user can reconstruct the top modelling solutions at atomic representation and continue with the analysis.

### Analysis

Analysis of the output model ensemble (Figure 1 step 8) entails the assessment of the sampling convergence and exhaustiveness, estimation of the sampling and model precision, quantification of restraint violation, and selection of representative models or model ensembles. For the assessment of exhaustiveness and precision, Assembline provides a command line interface to generate modelling output compatible with the imp-sampcon^20^ toolkit from IMP. The toolkit assesses the exhaustiveness based on four statistical criteria and estimates sampling precision as the highest precision at which the sampling can be considered exhaustive. For this, the output ensemble of good-scoring models is split randomly into two samples of approximately equal size. The first two tests assess the convergence of scores and similarities between the distribution of scores in the two model samples. The remaining two tests, performed upon structural clustering of the models from the two samples, evaluate the structural similarity of models in the samples by checking whether each cluster includes models from each sample proportionally to its size and if the modelled structures from the two samples are similar in each cluster. This analysis outputs several files describing the contents and metrics of each structural cluster and graphical plots summarizing the results of sampling precision assessment^20^.

### Comparison with other methods

Other integrative modelling software have been published and applied to a variety of systems. IMP^27^ and PMI^28^, which we use in our protocol, can on their own be used for integrative modelling. They offer the possibility to construct custom-made integrative modelling protocols based on multiple types of restraints, and defining the initial system architecture and its representation generically. However, a user of PMI or IMP would typically require Python programing expertise to modify and adapt the tutorial scripts to their specific modelling case. In contrast, our protocol brings an advantage of a versatile input configuration system consisting of a graphical interface and text configuration files that can be applied to a wider range of complexes without extra programming. This system enables configuring the input for even the most intricate complexes, for which structures of subunits or domains may be scattered across dozens of PDB files; parts of the complex might be subject to different restraints; or for which multiple symmetries might be present (as with the nuclear pore complexes and bacterial secretion systems). Our provided scripts can be applied to new systems without any modification. Despite the high-level interface, Assembline nevertheless offers the full functionality of IMP and PMI through a Python interface of Assembline. On top of IMP and PMI, our protocol implements its own algorithms for efficient sampling of the conformational space and provides additional modelling restraints. In particular, our fit library approach, which enables fast exploration of the conformational space, and the multi-step optimization algorithm are features not available in IMP or PMI. Thus, Assembline benefits from all features of IMP and PMI and extends them with novel and easily accessible functionalities.

Other tools, such as FoXS^37^, EMageFIT (from IMP) or MultiFit^38^, have been implemented using IMP as an underlying programming library. These tools, however, focus on specific applications and do not offer the full flexibility that Assembline or IMP and PMI provide. Our optimization approach is conceptually similar to MultiFit^38^, which also first performs discrete optimization of possible configurations followed by refinement. Some limitations of MultiFit^38^ that motivated us to develop a new protocol is the reliance of MultiFit^38^ on *a priori* density segmentation that defines sub-areas of the map for fitting, but which might exclude good fits before the optimization, less extensibility compared to IMP/PMI and Assembline, and no systematic input management system as included in Assembline. Thus, while MultiFit^38^ and Assembline could lead to similar results in some cases, Assembline offers more flexibility, and it is more universal, especially for large complexes.

HADDOCK^39^ and M3^40^ (which uses HADDOCK internally) offer integrative modelling algorithms specializing in higher-resolution modelling with a physical force-field applied to atomistic representations or a coarse-grained MARTINI^41^ representation. HADDOCK can be run either through a web-server or command line interface. Advantages of HADDOCK and M3 include full support of flexibility and the physical scoring function that can complement experimental restraints. Nevertheless, HADDOCK and M3, because of the high-resolution molecular representation, still cannot be applied to complexes as large as the nuclear pore complex, and do not provide a procedure for efficient sampling of the EM map at a large scale. Thus, HADDOCK and M3 could be used to refine smaller complexes modelled with Assembline or to refine selected interfaces in models of large complexes.

Another versatile package for integrative modelling is PyRy3D (http://genesilico.pl/pyry3d), which also offers accessible user interface and a variety of restraints. In comparison, Assembline enables more efficient sampling algorithms and, by integrating with IMP and PMI, provides more restraints and tools for analysis. In principle, ROSETTA software^42^ can also be used for integrative modelling and offers a vast spectrum of modelling algorithms and full support of structural flexibility. However, its versatility for integrative modelling applications depends on the development of additional customized protocols and thus a considerable level of modelling and programming expertise.

When only an EM map is available, Assembline can be used as a tool for simultaneous fitting of multiple components into EM maps. There are several other tools capable of producing models through single or multiple-fitting to medium- or low-resolution EM maps by flexible or rigid body fitting, for example: UCSF Chimera^26^, Situs^43^, Flex-EM^44^, MDFF^45^, γ-TEMPy^46^, CAMPARI^47^, iMODFIT^48^, MDFIT^49^, FOLD-EM^50^, or ATTRACT-EM^51^. For many EM-only applications and small complexes, these tools are sufficient and can be used instead of Assembline. Nevertheless, Assembline brings an advantage of a complete and versatile package that integrates all modelling steps, the possibility of restraints other than, and in addition to, EM, and an integrated pipeline for analysis of the final models.

In summary, Assembline brings broad usability and unique algorithmic advantages compared to other solutions and seamlessly integrates with complementary modelling programs.

### Limitations

Out-of-the-box, our protocol is designed for medium-to-low-resolution modelling, approximately worse than 4 Å resolution. Although it can be applied to fit subunit structures to high-resolution EM maps, other methods are more suitable for atomic-level *de novo* model building or flexible refinement of the resulting models. Currently, it does not support nucleic acids and offers only partial support for flexible deformation of structures during modelling through modelling of loops as flexible beads and elastic-network restraints between rigid bodies. We plan to address these two limitations in future versions of the protocol. As with most optimization algorithms, the protocol also relies on multiple parameters that have to be adjusted by the user, which we facilitate through appropriate guidelines and examples (see Supplementary Manual and Supplementary Tutorial 1,2).

### Expertise needed to implement the protocol

Users of Assembline should be familiar with the basic principles of structural modelling and structural analysis. Familiarity with Unix command line and executing command line programs is necessary, but the provided tutorials (Supplementary Tutorial 1, 2) assume only basic knowledge, and guide users through all steps with explanations. No programming skills are needed, but users should familiarize themselves with the syntax of the configuration files and graphical molecular visualization software, which is needed to set up the input and analyze the output. Finally, the ability to access and use a computer cluster is beneficial, especially for larger systems.

## Materials

### EQUIPMENT

#### Hardware

● Personal computer or computer cluster with minimum 50 GB of free disk space and 4 GB of RAM memory
● Internet connection to access the online versions of Assembline usage manual and tutorial material (optional), i.e., for *S. cerevisiae* NPC (ScNPC) and Elongator complex modelling

#### CRITICAL

For faster calculations, we advise to run Assembline on a workstation with as many processors as possible or on a computer cluster.

#### Input data sets

● Sequences of protein subunits to be modelled in the FASTA format
● Atomic protein structures (in the PDB format) that will be used to model the protein complex
● cryo-EM densities of the modelling target complex in the MRC format and/or XL-MS data in the Xlink Analyzer^25^ format

#### CRITICAL

Example data sets can be found in https://git.embl.de/rantos/scnpc_tutorial.git (ScNPC modelling material) and https://git.embl.de/kosinski/elongator_tutorial.git (Elongator complex modelling material). Other structural data types can be used through the standard IMP and PMI Python programming interface within a `custom_restraints()` function of the parameter file.

#### Example input files

● Input sequence FASTA file, PDB structures and cryo-EM maps used for integrative modelling of the cytoplasmic ring (CR) Y-complex from ScNPC^7^ (https://git.embl.de/rantos/scnpc_tutorial.git).
● Input sequence FASTA file, PDB structures, XL-MS data sets and negative-stain EM maps used for integrative modelling of the Elongator complex^29^ (https://git.embl.de/kosinski/elongator_tutorial.git).

### Software prerequisites

#### CRITICAL

Detailed instructions are provided for installation of all prerequisite software in the Assembline manual (Supplementary Manual and in the online version: https://assembline.readthedocs.io/en/latest/installation.html#).

● UNIX-based operating system (e.g., Linux, Ubuntu, CentOS)
● UCSF Chimera^26^ (free molecular visualization and analysis software)
● Xlink Analyzer^25^ plugin for UCSF Chimera^26^ (graphical interface for Assembline)
● Anaconda software (open-source distribution for Python and R for scientific programming, documentation: https://docs.anaconda.com/)
● Python programming language (version 3) distribution from Anaconda
● Assembline package (Anaconda-bundled package from `kosinskilab` channel in Anaconda)
● IMP^27^ (version 2.14 or newer, package from `salilab` channel in Anaconda)
● Scipy^52^, numpy^53^, scikit-learn^54^, matplotlib^55^ and pandas (http://doi.org/10.5281/zenodo.3715232) Python packages (packages from generic channel in Anaconda)
● pyRMSD^56^ package (package from `salilab` channel in Anaconda)
● hdbscan^57^ package (package from `conda-forge` channel in Anaconda)
● R programming language distribution from Anaconda
● fdrtool^34^, psych (https://cran.r-project.org/web/packages/psych/index.html), ggplot2 (https://cran.r-project.org/web/packages/ggplot2/index.html), tidyr^58^, data.table packages (packages from generic R channel in Anaconda)
● optional, Modeller^59^ package (comparative modelling software free for academic users)
● gnuplot package (http://www.gnuplot.info/)

#### CRITICAL

The software prerequisites along with the Assembline package that we describe in this work have been thoroughly tested in UNIX-based systems with `bash` terminal, therefore the protocol might not work as expected in OS X systems or other shell environments.

### Procedure

**Set up prior to modelling**

**Timing ∼2 h**

#### CRITICAL STEP

In order to proceed with the following steps, first make sure you have installed all prerequisites listed in the **Software prerequisites** section.

1. Activate the Assembline modelling virtual environment, where all software prerequisites have been installed, with the following command: `source activate Assembline` or, depending on your Anaconda setup: `conda activate Assembline`
2. Create a new directory for the modelling project, which will be referred to as the project directory, with the following example command for UNIX-based systems: `mkdir Elongator`

#### CRITICAL STEP

Input and configuration files for Assembline can be located in arbitrary locations on the disk, though it is recommended that they are stored in a single directory for easier management of the modelling project.

3. Create a single file (in the FASTA format) with all sequences of the target subunits (i.e., subunit is a protein of a target complex) and store it in the project directory. Example FASTA file with sequences for the Elongator complex is provided in https://git.embl.de/kosinski/elongator_tutorial.git (Elongator complex modelling material).
4. In the project directory create a new sub-directory and store all the structures of the target complex subunits in the PDB format. Note that the subunit chains can be organized in the PDB files in any way, e.g., a PDB file can contain single or multiple subunits, or extra proteins not used in modelling etc.

#### CRITICAL STEP

Make sure that the protein sequence and residue numbering in the PDB files correspond to the sequences in the FASTA file. Additionally, it is recommended to prepare the PDB files such that each PDB file would correspond to an anticipated rigid body (i.e., there is one-to-one mapping between the PDB files and the anticipated rigid bodies) that will be used to model the target complex.

5. In the project directory create a new sub-directory and store all the EM maps in the MRC format. These maps will be used for the definition of the EM restraints (read more about the EM restraints in Supplementary Manual).
6. Optionally, in the project directory create new directories and store other available data sets that will be used as input for integrative modelling. For example, create a new directory called `xlinks` and store XL-MS data sets in Xlink Analyzer format.

**Calculation of fit libraries**

**Timing ∼5 h**

#### CRITICAL STEP

The following steps for the calculation of fit libraries can be tested by retrieving and utilizing the fitting data and fitting parameters provided in https://git.embl.de/rantos/scnpc_tutorial.git (ScNPC modelling material) and https://git.embl.de/kosinski/elongator_tutorial.git (Elongator complex modelling material). Furthermore, the fit libraries will be used as “discrete restraints” (read more in Supplementary Manual) during the global optimization runs with Assembline.

7. Create and store in the project directory a parameter file that includes the paths to input PDB structures and EM maps to be used for fitting, the fitting parameters and options for execution of fitting on a computer cluster (recommended). An example parameter file is provided in https://git.embl.de/kosinski/elongator_tutorial.git (Elongator complex modelling material) and detailed explanations are provided in the Supplementary Manual.
8. Run the fitting of the specified input PDB structures in the experimental maps with the following command: `fit.py efitter_params.py` **TROUBLESHOOTING**
9. Upon completion of the fitting, start the analysis of the fit libraries by calculating the *p*-values of the individual fits with the following command: `genpval.py <FITTING directory name>` **TROUBLESHOOTING**
10. Optional, enter the directory with a prefix name that includes the fitting parameters (e.g., `results/search100000_metric_cam_inside0.3_radius500`) and generate the models (in the PDB format) of best scoring fits for visualization. For example, to generate top five fits from a fit library for each input structure: `genPDBs_many.py -n5 top5 */*/solutions.csv`

#### Global optimization

**Timing ∼1 h to days depending on the protein complex size and computational resources**

#### CRITICAL STEP

Note that the global optimization with Assembline should be run only upon successful generation of the fit libraries in the previous steps.

11. Open Xlink Analyzer graphical interface (Figure 2), which can be accessed as a plugin in UCSF Chimera, and create a project for the target complex, which will be used as the modelling configuration file (in the JSON format).

**Figure 2.**
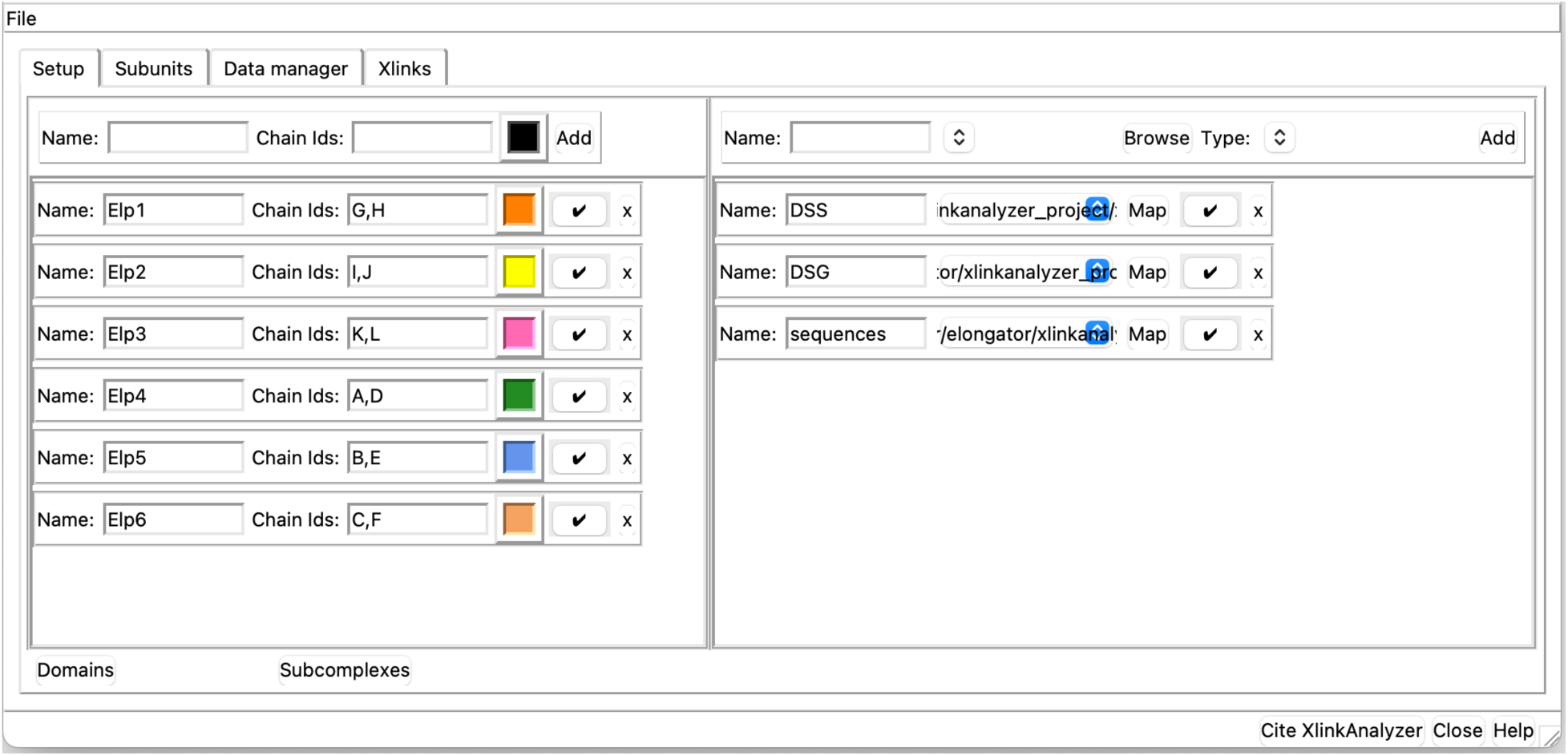
An example of Xlink Analyzer interface used for configuring the modelling project.

#### CRITICAL STEP

Read more on how to install Xlink Analyzer as a plugin for UCSF Chimera and how to create a project for the target complex from scratch in the tutorial of Xlink Analyzer described by Kosinski et al. (2015)^25^ (available at https://www.embl-hamburg.de/XlinkAnalyzer/XlinkAnalyzer.html) and the Elongator modelling tutorial (https://elongator-tutorial.readthedocs.io/en/latest/json_setup.html).

Subunits are defined in the panel on the left, while data is loaded and mapped to subunits in the panel on the right.

12. Add all subunits using the Xlink Analyzer interface.
13. Assign unique chain ID and color to every subunit.
14. Optionally, define domains within subunits, which can be used later for e.g., defining restraints specific to the domains.
15. Add sequences using the Setup panel in Xlink Analyzer and map the sequences to names of subunits using the Map button.
16. Optional, add available XL-MS data sets and map the crosslinked protein names to names of the subunits using the Map button.
17. Make a local copy of the Xlink Analyzer project file, e.g. copy the `XlinkAnalyzer_project.json` as `X_config.json.` The new `X_config.json` will be used as the configuration file for modelling, whereas the original file should be kept for analysis of models.
18. Open a text editor and edit `X_config.json` project file manually to add modelling directives, which include: definition of series (grouping of subunits and domains), symmetry information, definition of input PDB structures and rigid bodies, specification of fit libraries and definition of spatial restraints.

#### CRITICAL STEP

Due to the different complexity of every modelling project, and plethora of available options and combinations in setting up the configuration file, the detailed description of how to set up this configuration file along with explanations regarding the definition steps are provided in Supplementary Manual. We recommend text editors with syntax highlighting to edit the configuration file (in JSON format), for example, SublimeText (https://www.sublimetext.com/) and Atom (https://atom.io/). Example configuration files can be found in https://git.embl.de/rantos/scnpc_tutorial.git (ScNPC modelling material) and https://git.embl.de/kosinski/elongator_tutorial.git (Elongator complex modelling material).

19. Create a parameter file in Python language format (e.g., `params.py`), which defines the modelling protocol, scoring functions, output parameters and some restraints for the global optimization stage and save the file in the project directory.

#### CRITICAL STEP

Note that although the file is in the Python language, no programming skills are required. If the modelling is going to be performed on a computer cluster, this parameter file should also include cluster submission settings. Detailed description of how to set up this file are provided in Supplementary Manual. We recommend text editors with syntax highlighting to edit the configuration file (in JSON format), for example, SublimeText (https://www.sublimetext.com/) and Atom (https://atom.io/). Example configuration files can be found in https://git.embl.de/rantos/scnpc_tutorial.git (ScNPC modelling material) and https://git.embl.de/kosinski/elongator_tutorial.git (Elongator complex modelling material).

20. Using the Unix command line, navigate to the project directory and run the global optimization using the following example command, which will submit all runs to the computer cluster or to a standalone workstation in chunks of N models (i.e., parallel runs) according to the number of processors defined through `ntasks` parameter in the parameter file. The following command will submit 1,000 modelling jobs in the computer cluster queue or run them consecutively in chunks on a standalone workstation with each job leading to one model: `assembline.py --traj --models -o out --multi --start_idx 0 - -njobs 1000 X_config.json params.py &>log&` **TROUBLESHOOTING**

#### CRITICAL STEP

Note that the number of processors to be allocated for the modelling runs or cluster submission commands and templates should be specified *a priori* in the parameters file. Read more on how to customize the submission of multiple modelling runs according to the computational system architecture in Supplementary Manual. Additionally, it is suggested to perform a single test run before deploying multiple jobs to a computer cluster or even a local workstation with a command similar to the following example:

`assembline.py --traj --models --prefix 0000000 -o out config.json params.py`

21. Upon completion of the global optimization runs, enter the output directory, inspect the output log files to validate that the included information (e.g., the list of final parameter values used for modelling, the summary of the molecular system created, the scores of defined spatial restraints) match the expected output. **TROUBLESHOOTING**

#### CRITICAL STEP

In case that the output modelling logs do not include all the modelling parameters and restraints set prior to modelling then repeat the previous step after correcting the parameter file accordingly. Read more on how to edit the parameter file and how to evaluate the output logs in Supplementary Manual.

22. While in the output directory, generate reports with total scores and individual scoring terms for all modelling runs with the following command: `extract_scores.py` **TROUBLESHOOTING**

#### CRITICAL STEP

The `extract_scores.py` script will generate several files with lists of models and their respective scores. Make sure that the `all_scores.csv` file and `all_scores_sorted_uniq.csv` files were generated as they will be used as input for the analysis in the following steps.

23. Optionally, plot histograms of all total scores and scoring terms for all modelling runs from global optimization with the following command:

`plot_scores.R all_scores.csv`
24. Optionally, while in the output directory, generate the best scoring models from the global optimization run in the mmCIF format. For example, to generate the single best scoring model): `rebuild_atomic.py --project_dir <FULL path to the original project directory> --top 1 all_scores_sorted_uniq.csv` **TROUBLESHOOTING**

#### CRITICAL STEP

The `--project_dir` option is only necessary if relative paths were used in the JSON configuration file for global optimization. Also note that by default the `rebuild_atomic.py` script will generate only the parts of the models that correspond to the specified input rigid bodies, meaning that the flexible bead parts of the system (if any) will not be rebuilt. Read more on how to generate flexible beads of subunits, or even how to generate best-scoring models in formats other than mmCIF (e.g., the PDB format, although this format is not recommended for large systems) in Supplementary Manual.

25. Optional, visualize and inspect the modelling trajectory of the best-scoring models (or any other candidate model) by navigating and finding the corresponding trajectory file in `traj/` directory in the output folder from modelling and displaying it with UCSF Chimera. An example trajectory of the best scoring model from CR Y-complex (from the in-cell ScNPC model^7^) can be inspected in Supplementary Video 1. TROUBLESHOOTING
26. Optionally, while in the modelling output directory, run a quick sampling convergence test and inspect the convergence plots that will be stored in a PDF file called `convergence.pdf` with the following command (example command to include 20 trajectories from the modelling runs): `plot_convergence.R total_score_logs.txt 20`

#### CRITICAL STEP

If the plots clearly indicate no convergence, go back to step 19, increase the number of Monte Carlo steps and repeat global optimization.

27. While in the output modelling folder, create a file containing all subunits (and the residue indexes) that will be used to calculate localization probability densities (e.g., `density.txt`) during the sampling exhaustiveness assessment with `imp-sampcon exhaust` tool^20^ from IMP with the following command: `create_density_file.py --project_dir <PATH to the original project dir> config.json --by_rigid_body`

#### CRITICAL STEP

The output file (e.g., `density.txt`) from `create_density_file.py` script is required for the following steps of the sampling exhaustiveness analysis; therefore it has to be generated with respect to a very specific format. Read more on how to generate the `density.txt` file (as well as how to compile this file for complexes including symmetrical copies etc.) in Supplementary Manual. Example `density.txt` files can be found in https://git.embl.de/rantos/scnpc_tutorial.git (ScNPC modelling material) and https://git.embl.de/kosinski/elongator_tutorial.git (Elongator complex modelling material). Note that in case the modelled complex is homo-oligomeric, an extra file defining the symmetry is needed for the next step and can be generated using a command like the following example:

`create_symm_groups_file.py --project_dir <FULL path to project dir> config.json params.py`

28. Run the following command (`setup_analysis.py` script) which will automatically prepare the input files for the sampling exhaustiveness analysis with `imp-sampcon` tool^20^ from IMP based on the resulting integrative models: `setup_analysis.py -s <abs path to all_scores.csv file produced by extract_all_scores.py> \`

`-o <SPECIFIED output dir> \`
`-d <density.txt file generated in the previous step> \`
`-n <number of top scoring models to be analyzed, default is all models> \`
`-k <restraint score based on which to perform the analysis, default is total score>`
**TROUBLESHOOTING**

#### CRITICAL STEP

Read more about example commands and further options that can be used for the `setup_analysis.py` script in Supplementary Manual.

30. Enter the output analysis directory created by the previous step (i.e., output directory from `setup_analysis.py`) with the following command: `cd <ANALYSIS directory>`
31. Run the sampling exhaustiveness analysis with the `imp-sampcon exhaust` tool^20^ from IMP to assess the sampling performance of the modelling: `imp_sampcon exhaust -n <PREFIX for output files> \`

`--rmfA sample_A/sample_A_models.rmf3 \`
`--rmfB sample_B/sample_B_models.rmf3 \`
`--scoreA scoresA.txt --scoreB scoresB.txt \`
`-d <path to density.txt file>/density.txt \`
`-m <CALCULATOR selection> \`
`-c <NUMBER of processors to use> \`
`-gp \`
`-g <FLOAT with clustering threshold step> \`
`--ambiguity <symmetry group file if applicable)`

#### CRITICAL STEP

It is highly recommended to run the sampling exhaustiveness analysis on multiple-processors workstation or a computer cluster as some of the testing steps during this analysis are computationally demanding. Examples for cluster submission options for `imp-sampcon exhaust` are provided in Supplementary Manual and Supplementary Tutorial 1 and 2.

32. Inspect the four output plots and text files. If the sampling has not converged and sampling exhaustiveness has not been achieved, perform additional global optimization runs or adjust the modelling parameters, for example the number of Simulated Annealing steps.

**Recombination (optional)**

**Timing ∼1 h to days depending on the protein complex size and computational resources**

#### CRITICAL STEP

Note that the recombination step should only be run upon completion of the global optimization step.

32. While in the output directory of the global optimization run, use the following command to automatically generate the JSON format configuration file for this step: `setup_recombination.py \`

`--json <JSON file used for global optimization> \`
`--scores all_scores_uniq.csv \`
`-o <OUTPUT directory for the new fit libraries> \`
`--json_outfile <DESIRED name for JSON config for recombinations> \`
`--project_dir <ORIGINAL project dir> \`
`--score_thresh <SCORE threshold for selecting the models> \`
`--top <NUMBER of top scoring models to use for extracting the fit libraries>`
33. Navigate back to the main project directory and run a similar command to the following example, which will perform 1,000 modelling runs and store output in the global optimization output folder. The command will perform the modelling by recombinations with Assembline using the freshly generated JSON formatted configuration file (e.g. `config_recomb.json`) and modelling parameter file from global optimization: `assembline.py --traj --models -o out --multi --start_idx 0 - -njobs 1000 --prefix recomb config_recomb.json params.py &>log&` **TROUBLESHOOTING**

#### CRITICAL STEP

For recombination, usually the number of Simulated Annealing steps in `params.py` can be decreased for quicker calculations, because the used fit libraries are smaller. Read more on which settings are available and how to modify the parameter file generation procedure in Supplementary Manual.

34. Analyze the output from the modelling by rigid body recombinations with Assembline by following the exact analysis procedure described from step 21 up to step 31 (i.e., repeat steps 21-31). It is expected that this step generates additional good-scoring solutions. **TROUBLESHOOTING**

**Refinement**

**Timing ∼3 h to days depending on the protein complex size and computational resources**

#### CRITICAL STEP

Note that the refinement can be run even without prior global optimization (or recombination) if the input rigid bodies are already approximately fitted in the EM map. Also, if the results of the global optimization are satisfactory, this step can be skipped. However, it is recommended to run the refinement to further optimize the models. The refinement mode would be also used as the first and the only modelling step if no EM map is available. Read more regarding the refinement options in Supplementary Manual or inspect examples of refinement applications in Supplementary Tutorials 1 and 2.

35. While in the main project directory, run a command similar to the following example, which will automatically generate a generic configuration file (in the JSON format) based on the configuration and parameter files from global optimization: `gen_refinement_template.py --out_json refine_template.json - -params params.py --add_series X_config.json`
36. Open a text editor and edit the generated (from previous step) JSON configuration file manually to add modelling directives which might include: updated definitions of rigid bodies, restraints specific for the refinement, adjusted restraint weights.

#### CRITICAL STEP

The detailed description of how to set up this configuration file along with explanations regarding the definition steps are provided in Supplementary Manual. Example configuration files can be found in https://git.embl.de/rantos/scnpc_tutorial.git (ScNPC modelling material) and https://git.embl.de/kosinski/elongator_tutorial.git (Elongator complex modelling material).

37. In the main project directory, create and edit a copy of the global optimization parameter file in Python language format (if global optimization was not run beforehand then follow steps 11-19 to generate it from scratch). The final format of the file should be almost identical with the global optimization parameter file with the main differences being that the modelling protocol to be applied is `refinement` and the scoring function includes scoring terms derived from restraints specific to the refinement method of Assembline.

#### CRITICAL STEP

The detailed description of how to set up this file is provided in Supplementary Manual.

38. Prepare the top best scoring models from global optimization for refinement with a command similar to the following example (which will create a directory containing 100 folders with input data sets and configuration files for the top 100 models from global optimization): `setup_refine.py \`

`--top 100 \`
`--scores out/all_scores_uniq.csv \`
`--previous_json config.json \`
`--refine_json_template config_refine_template.json \`
`--refine_json_outname config_refine.json \`
`--previous_outdir out/\`
`--refine_outdir out/refinement`

#### CRITICAL STEP

In case the global optimization was not run beforehand and the refinement is applied to a pre-existing model, skip this step and specify in the configuration file the paths to the PDB files that will be used for refinement.

39. Run the refinement for all selected top-scoring models from global optimization (or single run in case global optimization was not run beforehand) with a series of commands similar to the following example (a `bash` loop that will apply refinement to the top 100 models from global optimization by running 10 refinement runs for each model, i.e. 1,000 runs in total): `for model_id in ‘ls --color=never out/refinement’;` 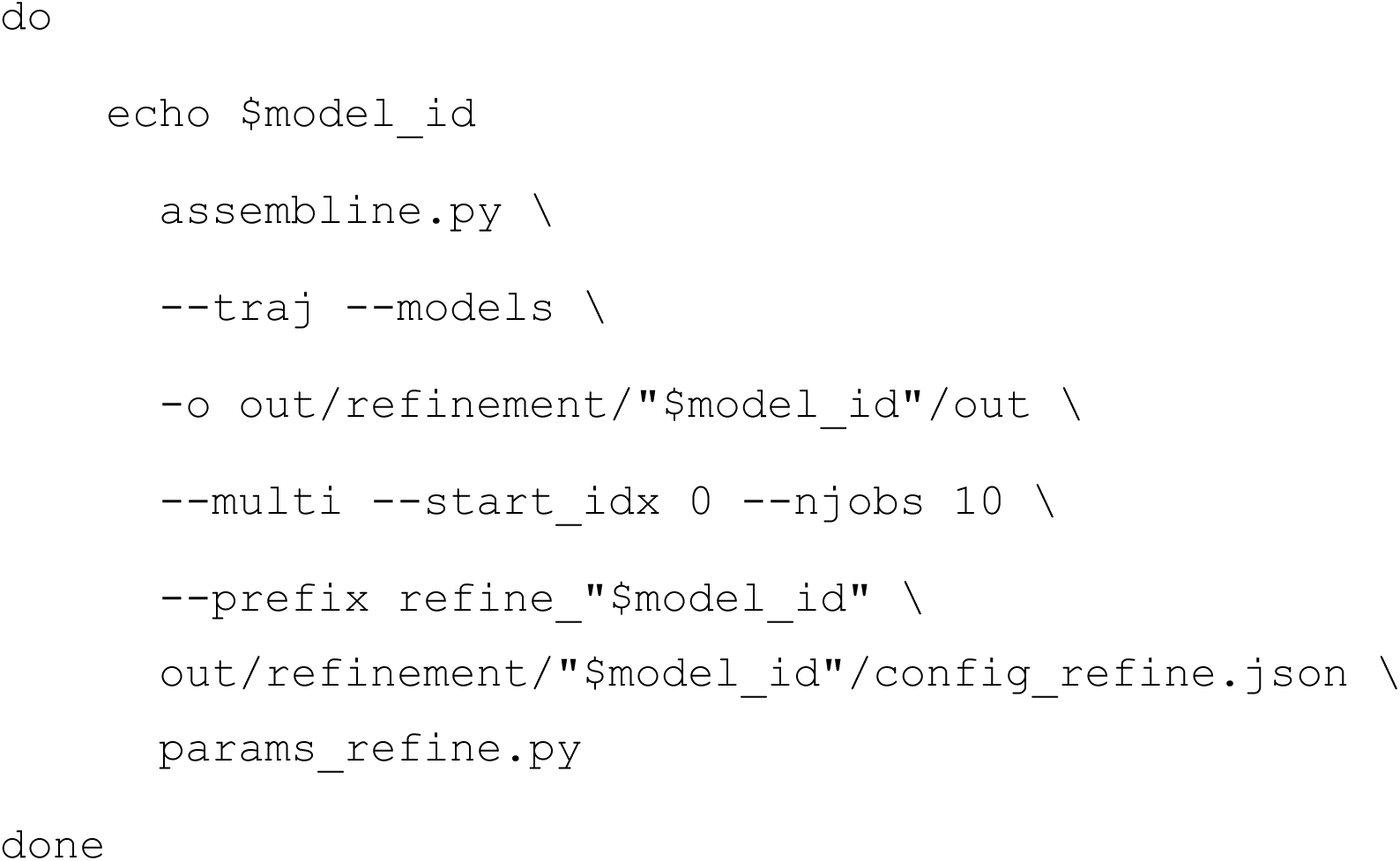 **TROUBLESHOOTING**

#### CRITICAL STEP

Note that the number of processors to be allocated for the modelling runs or cluster submission commands and templates should be specified *a priori* in the parameters file. Read more on how to customize the submission of multiple modelling runs according to the computational system architecture in Supplementary Manual. Additionally, it is suggested to perform a single test run before running multiple jobs with commands similar to the following example:

`model_id=‘ls --color=never out/refinement | head -n 1’ assembline.py \`

40. Analyze the output from the modelling by refinement by following the exact procedure described from step 21 up to step 31 (i.e., repeat steps 21-31), but now generating the scores using the command: `extract_scores.py --multi` 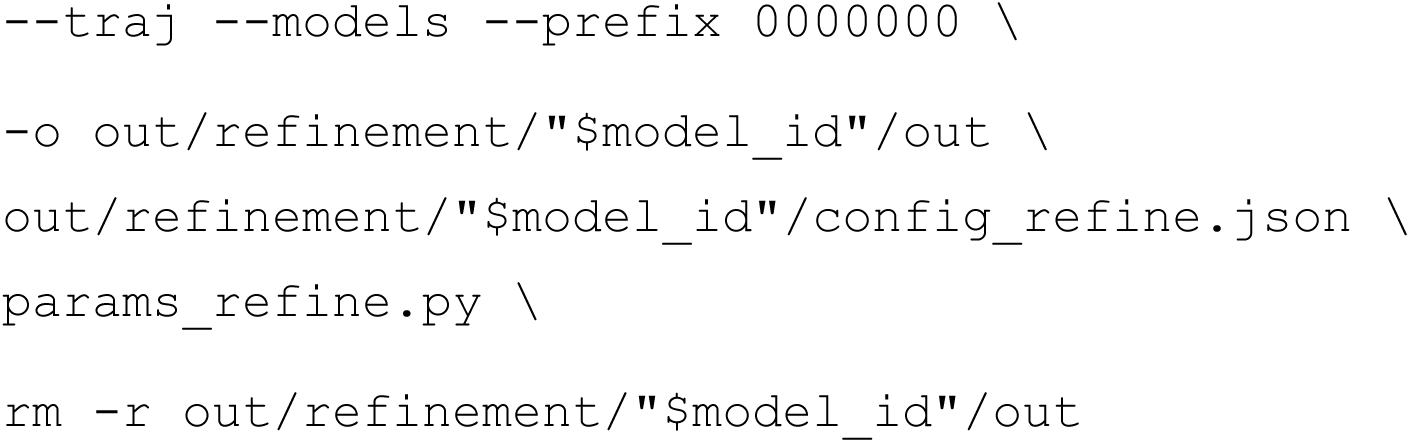 **TROUBLESHOOTING**

#### CRITICAL STEP

An example trajectory of the best scoring model (produced with refinement) from nuclear ring (NR) Y-complex (from the in-cell ScNPC model^7^) can be inspected in Supplementary Video 2.

### Troubleshooting

Troubleshooting advice can be found in Table 1.

### Timing

Steps 1-6, installation, set up of virtual environment for modelling and project directory architecture: ∼2 h per modelling project, assuming the input datasets are already prepared.

Steps 7-10, calculation of fit libraries: ∼1 h - 5 h, depending on number of input PDB structures to be fitted, number of EM maps to be used for fitting and allocated computational resources. The calculation of fit libraries for ScNPC takes roughly 3-5 h per input structure and for Elongator complex takes around 1-3 h per input structure.

Steps 11-31, modelling with global optimization and analysis pipeline: ∼1 h to days, depending on modelling parameters, system configuration settings, size of the complex, and allocated computational resources. A single global optimization run (using a single CPU core) for ScNPC takes around 1 h (approximately 20,000 runs to be performed) and for Elongator complex - 6 min (approximately 1,000 runs to be performed).

Steps 32-34, modelling with the recombination step and analysis pipeline: ∼1 h to days, depending on modelling parameters, system configuration settings, size of the complex, the number of selected models to be used for recombination, and allocated computational resources. A single recombination run (using a single CPU core) for Elongator complex takes around 4 min (approximately 1,000 runs to be performed).

Steps 35-40, modelling with refinement and analysis pipeline: ∼3 h to days, depending on modelling parameters, system configuration settings, size of the complex, the number of selected models to be used for refinement, and allocated computational resources. A single refinement run (using a single CPU core) for ScNPC takes around 3-4 h (approximately 20,000 runs to be performed) and for Elongator complex takes around 20 min (approximately 1,000 runs to be performed).

### Anticipated results

The post-processed output of Assembline is an ensemble of models, along with an analysis of model uncertainty. The models can be exported in either coarse-grained or atomic resolution as PDB or CIF files.

### Overview of integrative modelling output

The raw output of the integrative modelling stages in Assembline are models stored in bead representation, simple text files with rigid body transformations and the respective scores. Assembline provides a structural analysis toolkit with which any number of best scoring models can be converted to atomic representation and stored in the most common formats (e.g., PDB, CIF). For the production of the final atomic models, flexible loops can be rebuilt in a full atom representation with Modeller^59^ using the starting conformations of the loops derived from the bead representation.

As examples, in the following sections we describe how our protocol was applied in our recent study to model the nuclear pore complex from *S. cerevisiae*^7^ (Supplementary Tutorial 1). We also demonstrate the results of modelling of the Elongator complex from *S. cerevisiae*^29^, which was originally modelled with the older version of the protocol, but here is presented with the current version of Assembline and in a simplified setup suitable for an introductory tutorial (Supplementary Tutorial 2).

### Integrative modelling of ScNPC

The nuclear pore complexes (NPCs) are large macromolecular assemblies that fuse the nuclear envelope and facilitate nucleocytoplasmic transport^60^. They are built by around 30 different nucleoporins (Nups) present in multiple copies, which are organized in a triple-stacked ring conformation that forms a central transport channel^24, 60–62^. In a recent study^7^, we built models of ScNPCs based on in-cell cryo-ET data collected under wild-type and knock-out conditions.

The model of the wild-type ScNPC (Figure 3a) was constructed by integrating the data from cryo-ET maps of individual rings at approximately 25 Å resolution and biochemical data on protein-protein interactions and membrane-binding motifs^63^. Two of the three rings, the cytoplasmic (CR) and nuclear (NR) rings were modelled using the multi-step procedure of global optimization followed by refinement as described above (Figure 1). The inner ring (IR) model was constructed by immediately applying the refinement step to a previously published model of in vitro-purified ScNPC^64^.

**Figure 3.**
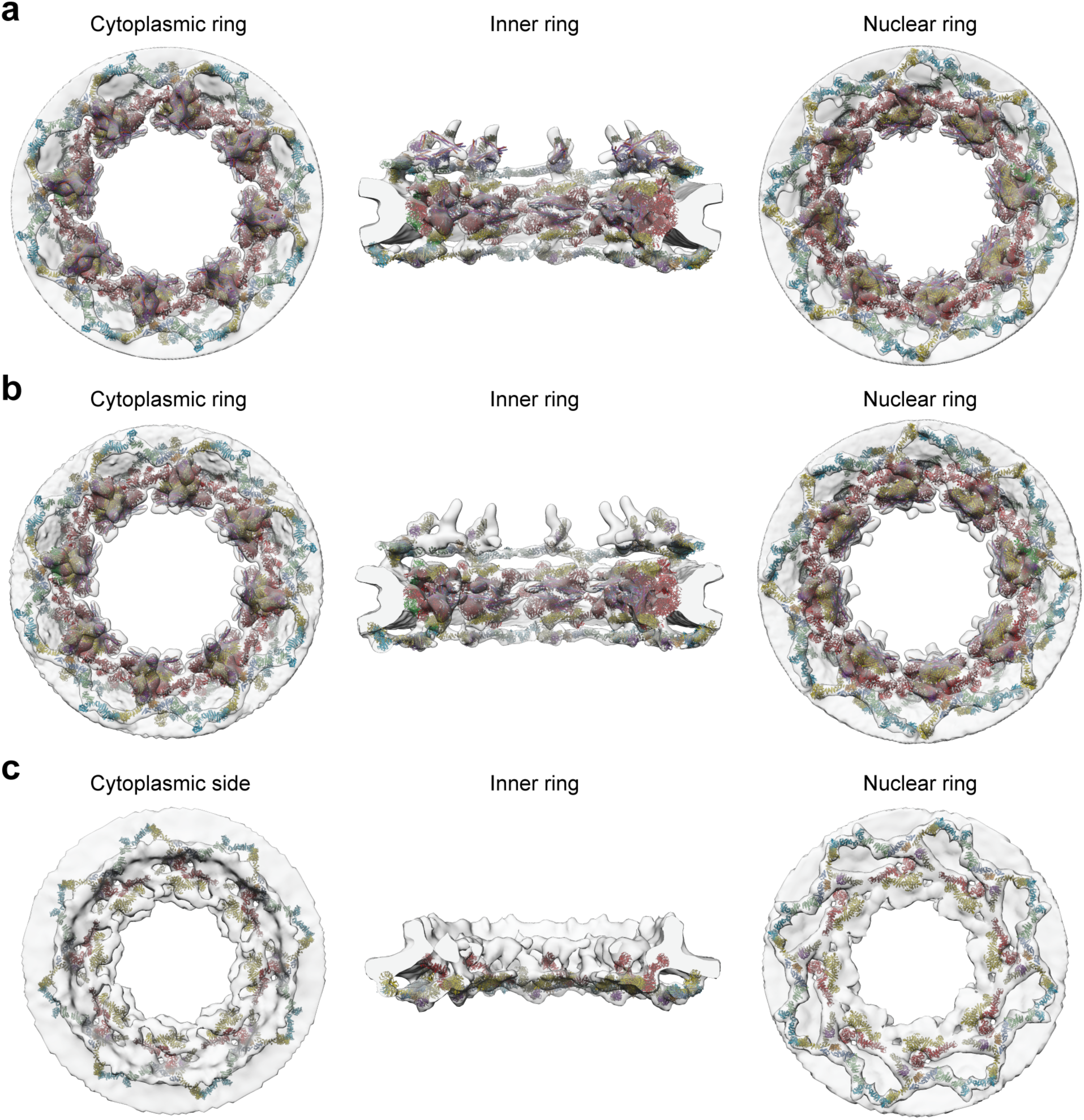
Integrative models of ScNPC. Overview of cytoplasmic ring (left), inner ring (middle) and nuclear ring (right) of **a,** the wild-type ScNPC, **b,** the ScNPC from cells with *nup116* gene knock-out (at 25 °C) and **c,** the ScNPC from cells with *nup116* gene knock-out (at 37 °C) where the cytoplasmic ring is entirely missing (left).

Two other cryo-ET datasets were collected under conditions where one of the subunits, Nup116 linking outer and the inner rings, was genetically knocked-out and cells were grown either at permissive (25 °C) or non-permissive (37 °C) temperature. The two conditions led to EM maps with approximately 25 Å and 50 Å resolution, respectively. Modelling was performed by first rigid-body fitting of the wild-type scaffold NPC rings to the knock-out EM maps and then optimizing the models using the refinement step directly, based on the EM maps and the interaction restraints. The refinement of the ScNPC model at the non-permissive temperature was challenging due to the low resolution of the EM map—the subunits were diverging from the initial structure and many different models were equally satisfying the restraints. As a troubleshooting solution, we applied an elastic network restraint available in Assembline, which enables the preservation of the interfaces between rigid bodies in the starting structure as long as the other restraints are not in conflict. The resulting *nup116* knock-out NPC model from cells grown under permissive temperature included the same scaffold complexes as the wild-type ScNPC and exhibited a very similar architecture, except for the missing density for Nup116 (Figure 3b). In the case of the *nup116* knock-out NPC model from cells grown under the non-permissive temperature, only the outer nuclear copies of the IR unit and the NR could be confidently included in the final model. Therefore, we hypothesized that our model represented a failed “inside-out” NPC assembly (Figure 3c).

Details and instructions in order to reproduce the wild-type and knock-out ScNPC modelling are provided in the Supplementary Tutorial 1 (and online at https://scnpc-tutorial.readthedocs.io/en/latest/).

### Integrative modelling of the Elongator complex

Elongator is a complex involved in tRNA modification^65^. In yeast, it contains six subunits, two copies each. In 2017, we published the integrative model of the yeast Elongator based on negative stain EM map at resolution of 27 Å and XL-MS data^29^. For the purpose of this protocol article, we repeated the modelling of the Elp123 subcomplex of Elongator by applying the updated procedure presented in this work.

The Elongator modelling case contained three subunits, each in two copies. The crystal structures and homology models of entire proteins or individual domains were used as input, grouped in nine rigid bodies (four rigid bodies for each of the two asymmetric units and one rigid body encompassing two copies of the same subunit) (Figure 4a and Supplementary Tutorial 2). The two-fold symmetry was applied as a constraint. From the calculation of fit libraries, between 200 and 500 fits were obtained for each rigid body. In the global optimization step, 1,000 models were generated. At this stage, the models already converged to a specific architecture (Figure 4b) leading to eight clusters at the sampling precision of around 10 Å and individual cluster (or model) precision between 7 Å and 20 Å. The top-scoring models represented very good fits to the EM map and only slightly violated the crosslink distance of 30 Å (Figure 4b). Since the sampling was exhaustive and at high-sampling precision, the recombination stage did not need to be performed. The top 100 unique models were refined, performing ten refinement runs for each of the starting models, yielding 1,000 refined models. Two loop regions harboring crosslinks were treated flexibly during refinement as chain of C*_α_* chain. The sampling precision obtained was approximately 20 Å and four clusters were obtained with precision of individual clusters between 10 Å and 20 Å. The top scoring model from refinement belonged to the largest cluster and satisfied all crosslinks (Figure 4c).

**Figure 4.**
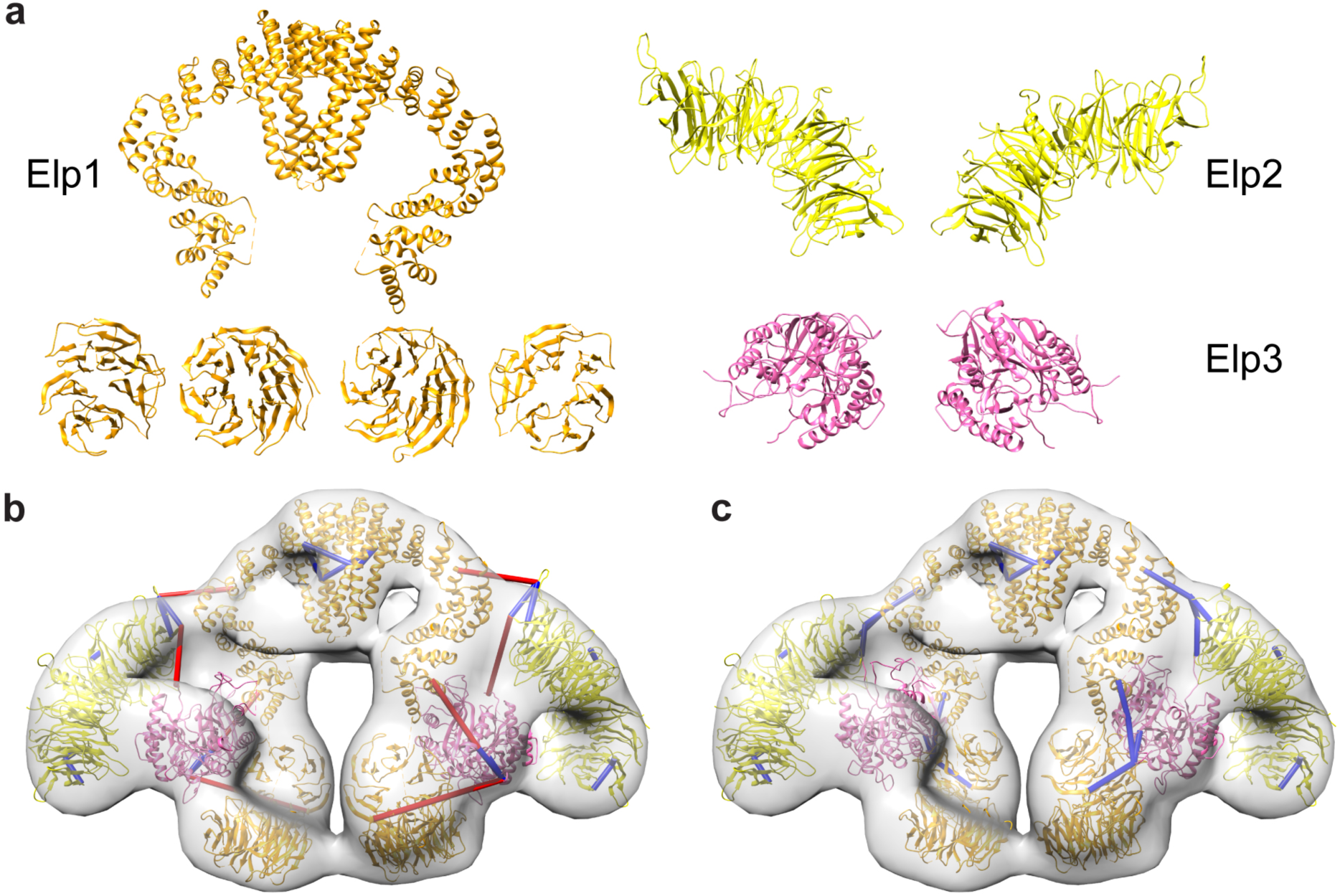
Integrative modelling of Elongator complex. **a**, Input structures (rigid bodies) used for modelling. **b,** The top-scoring model after global optimization within the EM map (gray transparent surface) and crosslinks indicated as blue and red bars (with red color indicating the crosslinks exceeding the expected distance of 30 Å, and blue - satisfying this distance). **c,** The top-scoring model after the refinement satisfies all crosslinks.

One challenge that we encountered in this case, which is common for negative stain EM maps, is that the calculated fitting libraries contained a very low number of alternative fits when generated with default options. This is likely due to high-density in few regions of the map, which are not reflecting the internal structure of the complex, but are an artifact of the negative stain method. The fit libraries are generated with UCSF Chimera by placing input structures in random positions in the map and then optimizing their local fit according to the EM cross-correlation score. If high-density regions are present, the optimization would shift the fitted structures to those regions, as they would give higher scores, resulting in most of the fits falling in the same locations. Thus, in the subsequent global optimization the models contained many clashes and did not lead to plausible models. To overcome this, we generated fit libraries with decreased clustering thresholds and less optimization steps, which led to a wider spectrum of fits and good scoring models from the global optimization.

The 3.7 Å cryo-EM structure of Elongator^30^, published in 2019, confirmed the model (Figure 5). The entire architecture was predicted correctly, including not only the localization of the domains but also their orientations. Although some features could be predicted only approximately, such as the exact orientation of the Elp3 subunit and the N-terminal domains of Elp1, functional sites were correctly localized relative to other subunits. This demonstrates that integrative modelling can generate models that enable functional interpretation of the structure.

**Figure 5.**
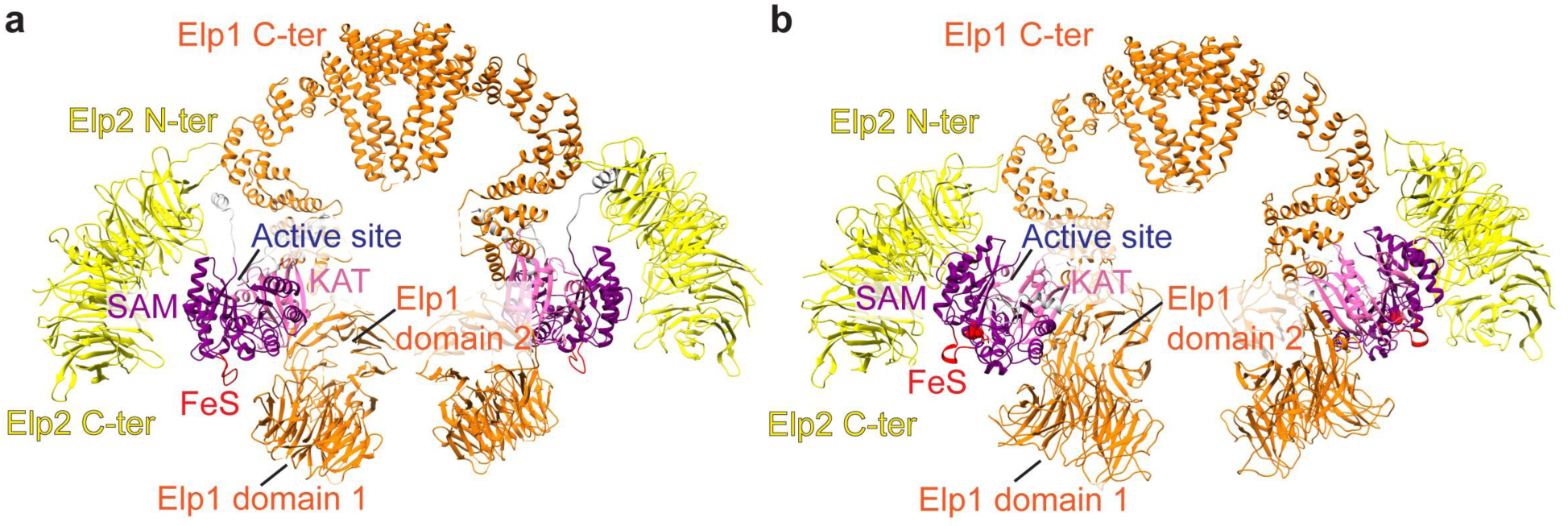
Comparison of the integrative model. **(a) and the high-resolution cryo-EM structure (b) of the Elp123 subcomplex of Elongator.** The subunits, domains, and functional sites are indicated for comparison. C-ter – the C-terminus, N-ter – N-terminus.

Details and instructions in order to reproduce the Elongator modelling are provided in Supplementary Tutorial 2 (and online at https://elongator-tutorial.readthedocs.io/en/latest/).

## Acknowledgements

We thank co-authors of the publications in which the Assembline protocol has been applied. We are grateful to Daniel Ziemianowicz and Karol Kaszuba for comments on the manuscript and the protocol, and Agnieszka Obarska for feedback on the protocol. The work has been supported by the Federal Ministry of Education and Research of Germany (FKZ 031L0100)

## Author information

### Contributions

V.R., K.K., and J.K. developed the protocol. V.R and J.K wrote the manuscript.

## Supplementary information

### Supplementary Manual

**Assembline installation and usage manual.** Detailed manual for Assembline protocol (online version: https://assembline.readthedocs.io/en/latest/<u>#</u>), which includes step-by-step explanations and suggestions on how to set up and run integrative modelling for any protein complex.

### Supplementary Tutorial 1

**ScNPC modelling tutorial.** Step-by-step tutorial including explanations for reproducing the ScNPC modelling based on in-cell cryo-ET maps as described in Allegretti et al.^7^ (online version: https://scnpc-tutorial.readthedocs.io/en/latest/<u>#</u>).

### Supplementary Tutorial 2

**Elongator complex modelling tutorial.** Step-by-step tutorial including explanations and suggestions for reproducing the Elongator complex modelling based on negative stain EM map and XL-MS data as described in Dauden et al.^29^ (online version: https://elongator-tutorial.readthedocs.io/en/latest/index.html<u>#</u>).

### Supplementary Video 1

**CR Y-complex trajectory from global optimization.** Animation video depicting the trajectory of the best scoring CR Y-complex model of the wild-type in-cell ScNPC^7^ produced with global optimization step. The model is shown in the coarse-grained representation inside the respective EM map.

### Supplementary Video 2

**NR Y-complex trajectory from refinement.** Animation video depicting the trajectory of the best scoring NR Y-complex model of the in-cell wild-type ScNPC^7^ produced with refinement step. The model (and the symmetrical copies) is shown in the coarse-grained representation inside the respective EM map.

